# Network theory predicts ecosystem robustness across environmental conditions

**DOI:** 10.1101/2025.01.29.635264

**Authors:** Germain Agazzi, Camille Carpentier, Olivia Bleeckx, Frédérik De Laender

## Abstract

Theory posits that a network’s link-species relationship predicts how changes in species richness *S* lead to changes in the number of links *L* between species. This relationship quantifies resistance to secondary extinctions and therefore gives pivotal information on ecosystem robustness. However, changes in *S* often coincide with environmental shifts, which can lead to unexpected impacts on *L*. In this paper, we constructed link-species relationships from 18 ecosystems using 1081 observations collected across different environmental conditions. We found that environmental noise (unspecified spatio-temporal variation) and environmental gradients (directional change) profoundly affected ecological network size (*S* and *L*), community composition, and induced network rewiring, with a Novotny’s rewiring index up to 0.47. Yet, we found the log(*L*) ∼ log(*S*) relationship to be remarkably constant across environmental conditions. Only in two of the 18 ecosystems did we find changes in environmental conditions to shift the log(*L*) ∼ log(*S*) relationship down, implying an overall drop in *L* but not in how species loss affects *L*. Our results show that network theory predicts ecosystem robustness across environmental gradients, which is encouraging for conservation.

## Introduction

Ecosystems are both complex [1–3] and of key importance to humans [4–6]. Understanding their response to global change drivers is therefore a priority for science and society [7–10]. Ecological networks are fundamental descriptors of ecosystem size and species interactions, and are therefore commonly used to understand ecosystem dynamics [4, 11]. These networks contain species as nodes and interspecific interactions as links (edges) [11–13].

The simplest way to define an ecological network is by its number of species (*S*) and number of links (*L*) [8, 13, 14]. As every species must interact with other species if it is to belong to the network, this relationship is necessarily positive, although the exact shape varies [15–20]. Historically, scientists proposed that a universal link-species relationship existed, along which every ecosystem could be located [19, 20]. However, more recently, the link-species relationship has been considered as network-specific [12, 15]. This line of thought postulates that the link-species relationship provides information on how a change in species richness within the network affects the number of links. The link-species relationship therefore readily connects to ecosystem response to species introductions [21] and extinctions, during network assembly and disassembly [9, 22–24].

A commonly used shape of the link (*L*)- species (*S*) relationship is the power law [15, 16], which turns into a linear function after logarithmic transformation of *L* and *S*:

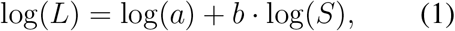

where the network-specific viewpoint imposes *a* and *b* to vary across networks. How changes of species richness *S* would therefore impact a network’s structure (*L*) is then predicted by the slope *b* [15], which is therefore inherently linked to ecosystem robustness [4, 8, 13, 15], one of the most studied ecological network properties [4, 7, 8].

Species losses or gains rarely occur at random but are driven by environmental changes [25]. These changes can lead to changes in network structure that surpass those induced by the mere change in species richness. That is because different environmental conditions can favor different species, which would lead to important shifts in community composition [25, 26]. Different species can imply different interaction schemes, but also when composition remains constant across environments species can change their interaction partners [5, 27, 28]. This implies that the parameters of Eq. 1 can change with environmental conditions: either the intercept (Fig. 1 A), or the slope (Fig. 1 B), or both. Alternatively, as proposed by Warren in 1990, the link-species relationship does not change with environmental conditions [29]. Thus, changes in species richness *S* would lead to predictable changes of *L*.

**Figure 1:**
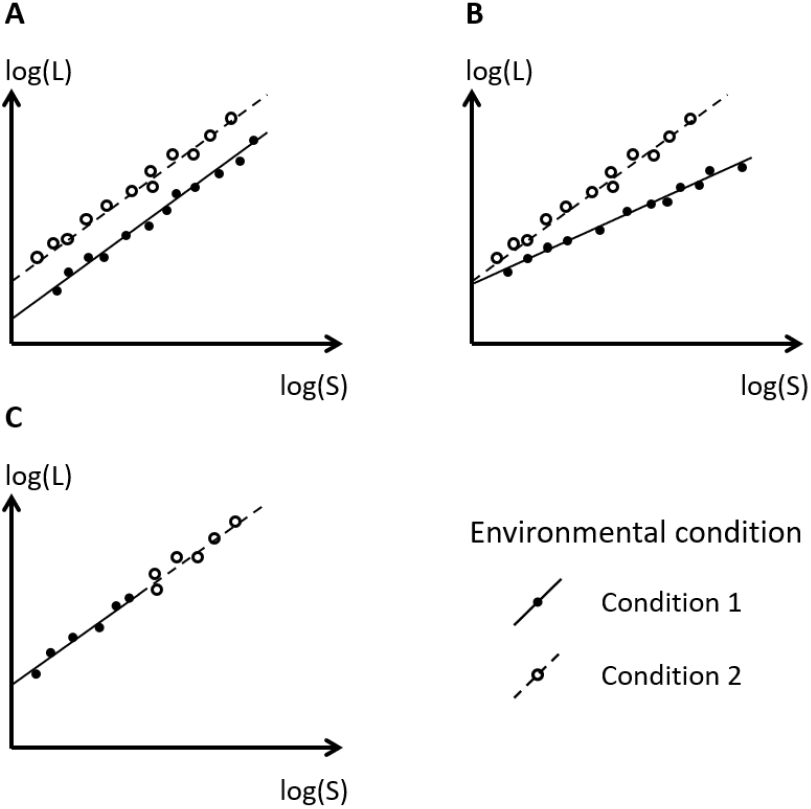
Possible impacts of changing environmental conditions on the log(*L*) ∼ log(*S*) relationship. (A) Conditions affect the intercept; (B) Conditions affect the slope, (C) Warren’s hypothesis: Conditions do not affect the relationship and effects on *S* lead to predictable effects on *L*. Combinations of cases A-C are possible but not shown here.

In this paper, we collected ecological network data from 18 ecosystems sampled at different environmental conditions. We considered two cases of changing conditions: environmental gradients (caused by ecosystem manipulation) and environmental noise (caused by spatiotemporal variation of unspecified environmental variables). We found that environmental gradients impacted the number of species and links (*S* and *L*), and induced compositional shifts and rewiring. However, despite these profound changes, the log(*L*) ∼ log(*S*) relationships did not change, supporting Warren’s hypothesis. This result implies that an ecological network’s link-species relationship predicts ecosystem robustness to biodiversity change, regardless of the environmental driver that is responsible for this change. These findings have deep implications for ecosystem monitoring and the assessment of ecosystem robustness.

## Results

We collated 1081 observations of the number of species *S* and links *L* across 18 ecosystems and a variety of environmental conditions (see supplementary material A) [30–41]. Changes in environmental conditions within an ecosystem either represented unspecified environmental noise caused by spatiotemporal variation [33, 35, 37–39, 41], or clearly defined environmental gradients caused by human ecosystem manipulation [32, 33, 36, 38]), or by specific environmental variables such as altitude [35], sun exposition [37], soil composition [39], season [40]), or succession state [41].

In the environmental noise category, only temporal variation significantly impacted *S* and/or *L* (see supplementary material B). Spatial variation never affected either net-work property. Environmental gradients had stronger effects on *S* and *L*. Specifically, sun exposure, mowing methods, buffalo presence, and soil composition affected both *S* and *L*. Fire frequencies affected only *S*, while successional states affected only *L*. Thus, some sources of environmental noise, and most environmental gradients, impacted ecological network size by changing *S* and/or *L* (Fig. 2).

**Figure 2:**
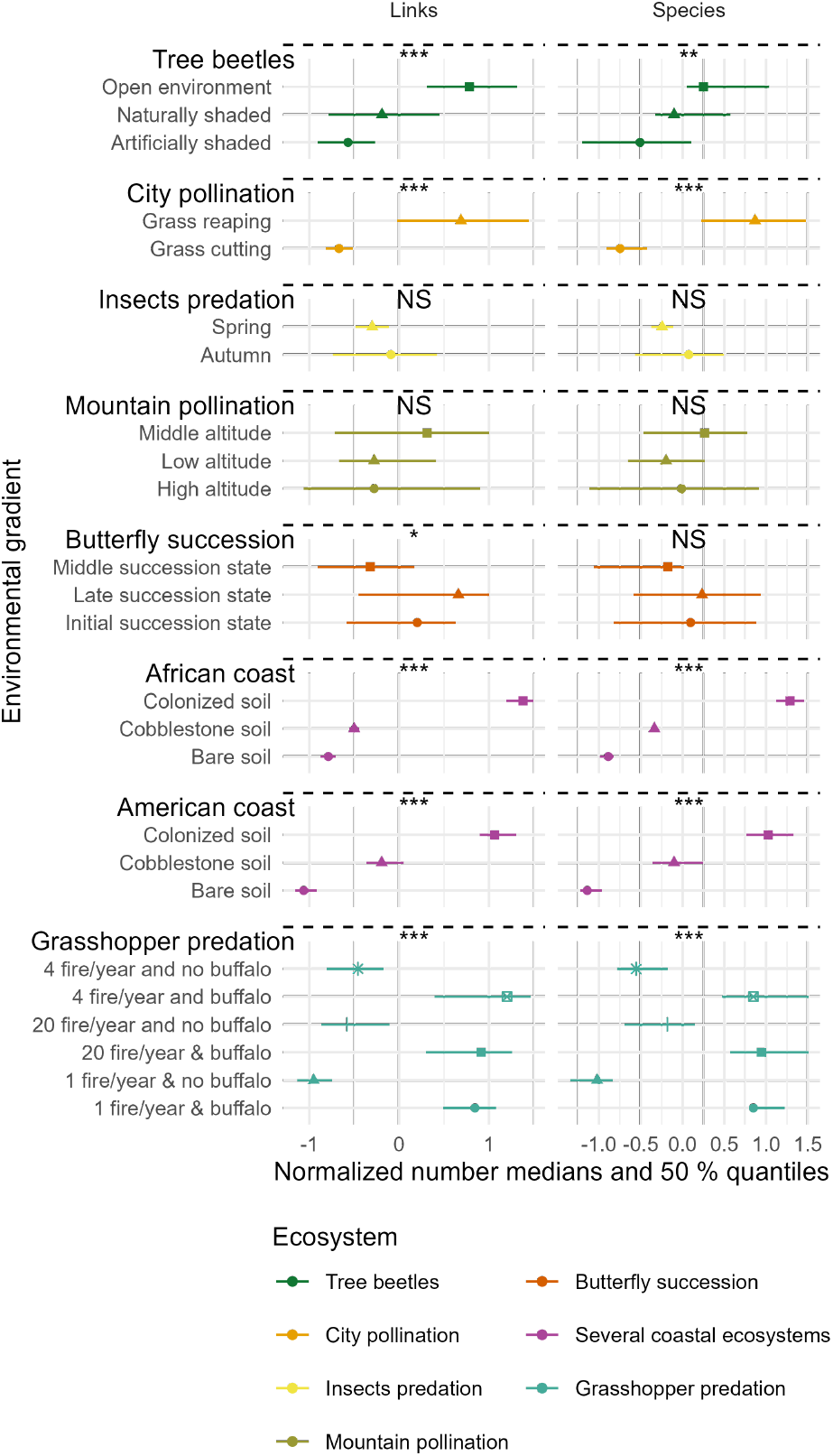
Environmental gradients affect the number of species *S* and links *L*. The dots represent the median of normalized numbers of species *S* and links *L* and the lines correspond to 50% of the variation. The kinds of environmental noise are defined in the supplementary material B.

Environmental conditions changes (gradients and noise) caused strong compositional shifts and rewired interactions of those species that persisted through environmental change (Fig. 3). Specifically, we tested how community composition changed with environmental conditions, for each ecosystem, using a Jaccard dissimilarity index. Values for this index vary between 0 (same composition in both environmental conditions) and 1 (environmental conditions have no species in common) [42]. We observed a median of 0.652 with middle 50% of data between 0.500 and 0.784. To measure if species that were present in both environmental conditions changed their interaction partners, we calculated a rewiring index [5, 27, 28, 43]. Using Novotny’s method, we found rewiring in 15 out of 18 ecosystems (Fig. 3).

**Figure 3:**
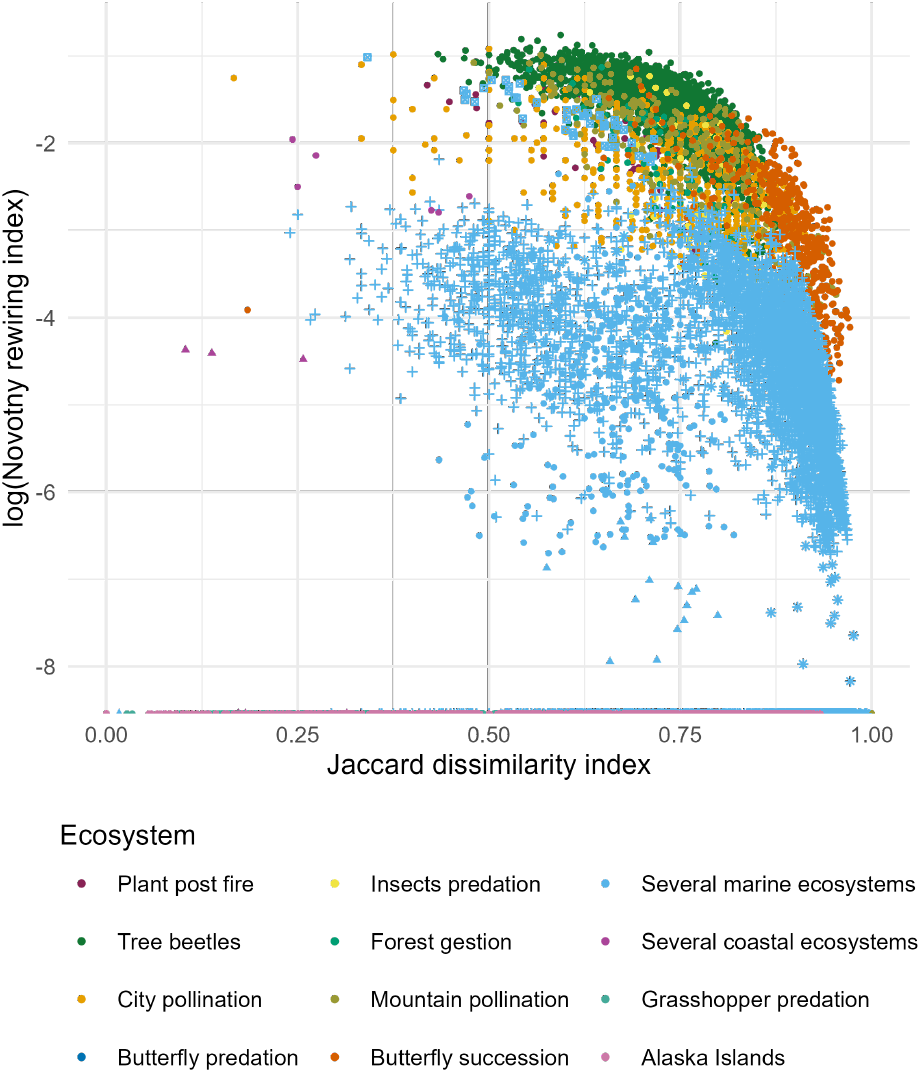
Environmental conditions changes cause compositional shifts and network rewiring. For each ecosystem, we compared community composition and network structure between two network observations and did so for all possible pairs of observations. For composition, we used Jaccard’s index; for network structure, we used the logarithm of Novotny’s rewiring index. Three ecosystems have a rewiring index of 0 (plotted along the horizontal axis). Details of all the calculations are in supplementary material C and details of the Jaccard index are in supplementary material D.

The power law was a good fit for all link-species relationships (Fig. 4). Regression analysis revealed that different ecosystems had different link-species relationships log(*L*) ∼ log(*S*) with significant differences in intercept and slopes (Fig. 4). We then carried out a model selection procedure that compared a model with log(*S*) and environmental condition (and their interaction) as predictors of log(*L*), against one that only contained log(*S*). We found no effects of environmental noise on log(*L*) ∼ log(*S*): changes in log(*S*) predicted changes in log(*L*) for all ecosystems (see supplementary material B). We also did not find significant impacts of environmental gradients on the slope of log(*L*) ∼ log(*S*), indicating that species loss or gain will have consistent effects on log(*L*), regardless of the environmental gradient causing it. We only found significant impacts on the intercept of log(*L*) ∼ log(*S*) for the case of sun exposition, buffalo presence, and fire frequencies (Fig. 5). Note that these environmental gradients also significantly affected *S* and/or *L* (Fig. 2). Across all selected models, all observations fell within the 95% prediction area. Moreover, models using only log(*S*) as a predictor of log(*L*) had a median *R*^2^ of 0.93, with 50% of the values ranging between 0.896 and 0.957. All *R*^2^ values exceeded 0.84, except one at 0.7, while the highest reached 0.996.

**Figure 4:**
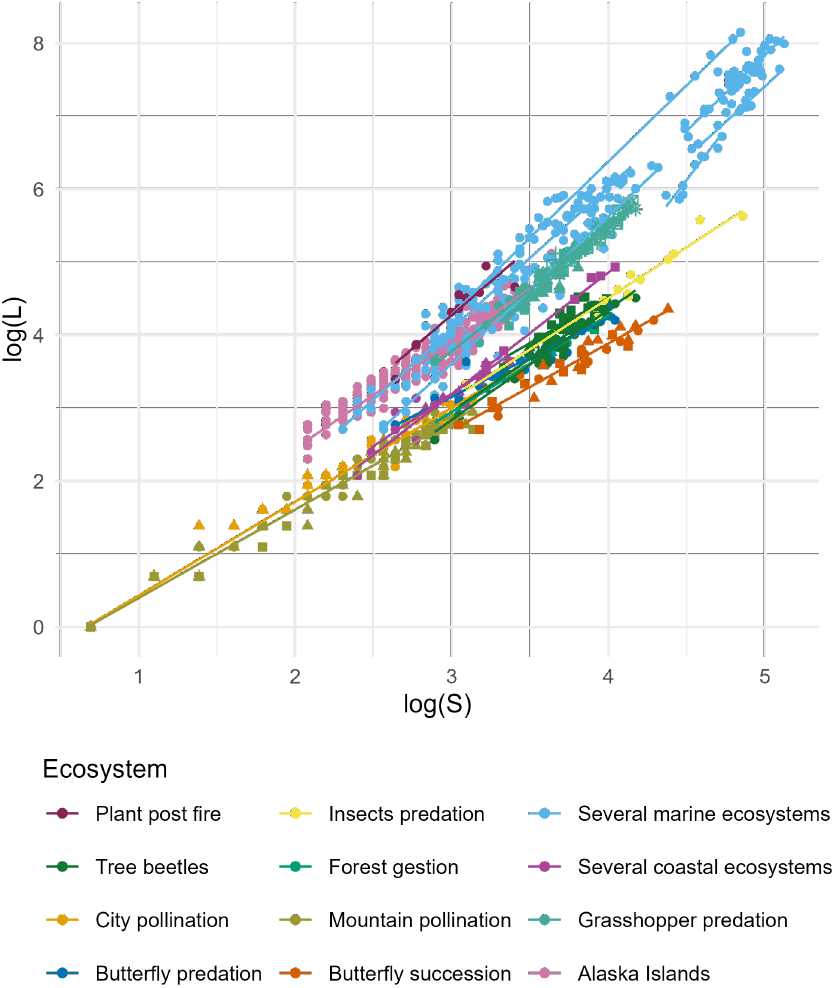
Different ecosystems have different relationships between species richness *S* and the number of links *L*, log(*L*) ∼ log(*S*). Colors represent ecosystems and shapes represent different environmental conditions (details in fig 2 and 5, and in supplementary material A). Lines represent log(*L*) ∼ log(*S*) relationships fitted to the data.

**Figure 5:**
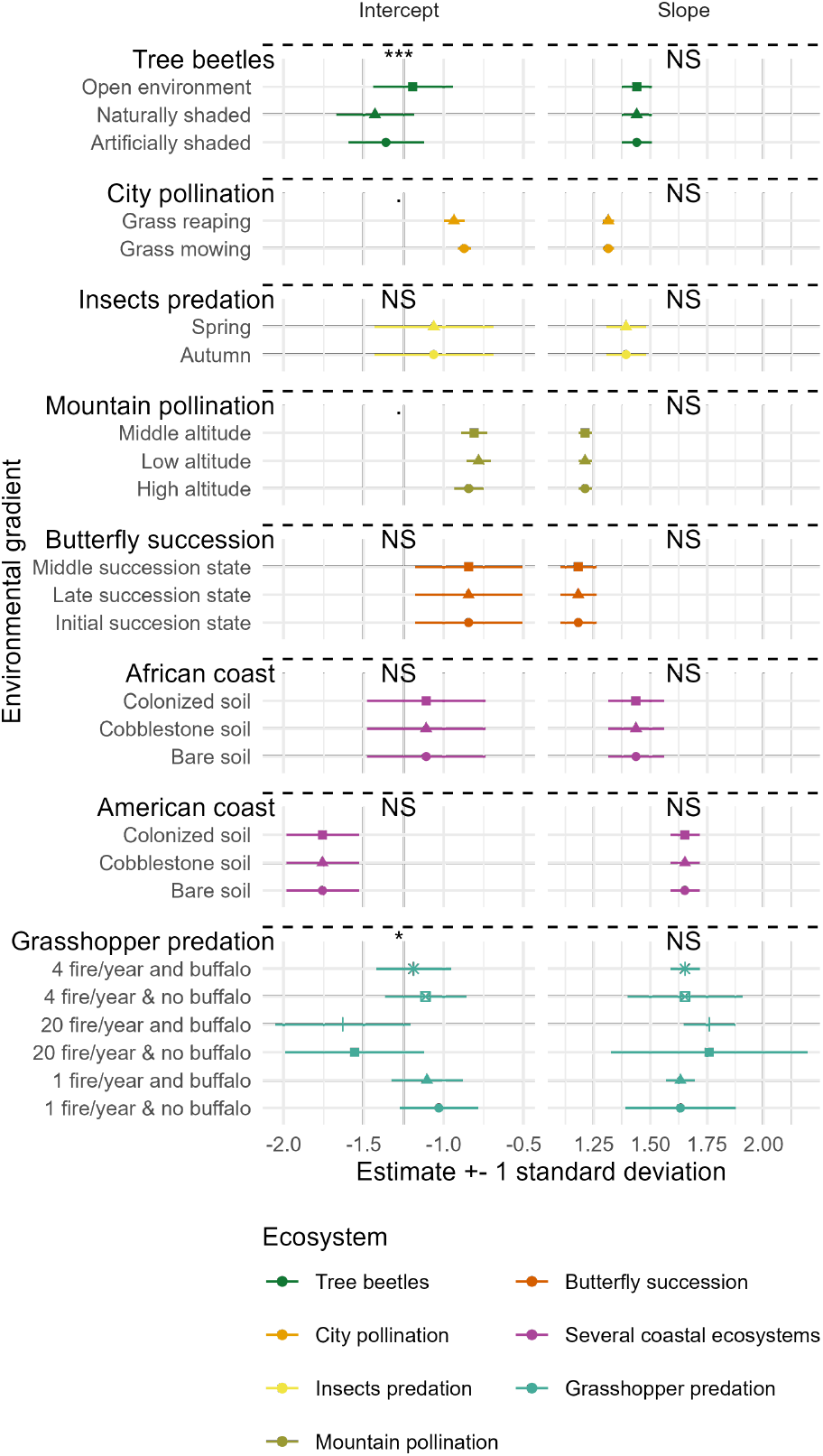
Environmental gradients rarely affected the intercept of the link-species relationship log(*L*) ∼ log(*S*), and never its slope. Dots represent the estimation by mixed or non-mixed linear models while the lines cover one standard deviation on this estimate. Environmental noise never affected link-species relationships (details are in supplementary material B).

## Discussion

We found that different environmental conditions lead to different network sizes (number of species *S*, and links *L*), different network compositions, and induced network rewiring, where species changed interaction partners as environmental conditions changed. This was most evident along environmental gradients. We did, however, find no evidence that the slope of the log(*L*) ∼ log(*S*) relationship depends on environmental conditions. Given that the slope of this relationship dic-tates ecosystem robustness, our findings imply that robustness is an inherent ecosystem property that predicts the functional consequences of biodiversity change, regardless of the environmental driver(s) of this change.

At a fundamental level, our results confirm that log(*L*) ∼ log(*S*) relationships are ecosystem-specific, as they have significantly different slopes and intercepts (Fig. 4) [15, 19, 20, 29]. If these differences among ecosystems are due to biological processes, or to different methodologies used by different authors is unknown [44]. Nevertheless, the inferred slopes vary between 1 and 2, confirming variation across ecosystems proposed before [12,15,45]. Combining this result with the rarity of environmental effects on the log(*L*) ∼ log(*S*) relationship (Fig. 5) suggests that ecosystem effects are stronger than environmental effects.

The invariance of the link-species relationship with environmental conditions suggests that other ecological network properties are invariant as well. For example, a fundamental network property is the connectance (*C*) which is the probability for a pair of species to interact and is measured as the proportion of potential links that are realized 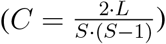 [4, 8, 13, 14, 46, 47]. As the number of links is *L* = *a* · *S*^*b*^ [15, 16, 19], connectance at high *S* can be approximated as *C* ≈ 2 · *a* · *S*^*b−*2^ with *b* predicted between 1 and 2 [12, 15, 45]. Thus, our results do show that some of the investigated environmental drivers may impact connectance by impacting *a*, as proposed before [48, 49]. However, even when neither *a* nor *b* changes, connectance will change following severe species loss (leading to low *S*), because of the inverse power-law shape of the *C* ∼ *S* relationship (Fig. 6) [4, 17, 30, 47, 50, 51].

**Figure 6:**
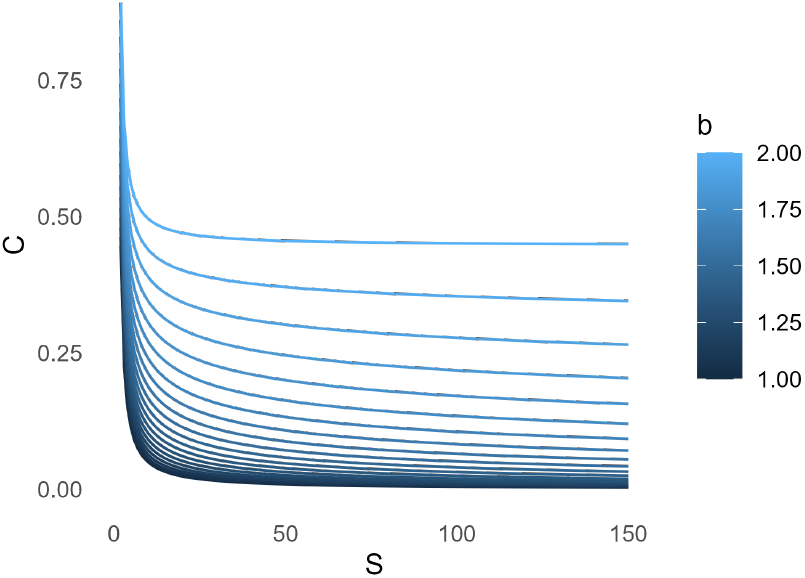
Connectance of an ecological network with constant *a* and *b*. The *b* value changes according to the legend and the *a* value is *e*^*−*1.5^ which matches the links-species relationships found here.

We found that most environmental gradients significantly affected the number of species and/or links (Fig. 2). Furthermore, we found that species compositions were profoundly impacted, with most of Jaccard dissimilarity indexes over 0.5 and that some species changed their interactions, causing rewiring of the network. Still, for 16 out of 18 of our link-species relationship, the slope of the log(*L*) ∼ log(*S*) relationship did not change. These results show how compositional turnover, as often observed [5, 40, 52–54], need not change the fundamental properties of ecological networks. This result corresponds to the hypothesis proposed by Warren that every ecological network has its link-species relationship, along which *S* and *L* change as environmental conditions change [29]. This result suggests two mechanisms.

First, it is possible that environmental drivers did not affect species based on how they were embedded in the network (e.g. removal of a keystone species at the start of an environmental gradient [55]). Second, it could mean that the remaining species adjusted their position in the network, in response to the new condition. Our results on rewiring suggest that the second option is plausible, but the contribution of the two proposed mechanisms is at present unknown.

Our results on rewiring highlight that ecological networks are highly dynamic, despite apparent topological consistency [22, 56] during assembly [41] or seasonal changes [40]. Nevertheless, we did not observe rewiring in all ecosystems. One explanation for this can be that some methods tend to introduce a bias towards no rewiring. In [44], for example, the species interactions were not directly observed but inferred from the literature. In such cases, rewiring cannot be observed, because species interactions only depend on the presence/absence of species.

The implications for ecosystem robustness require careful interpretation. Robustness describes the sensitivity of an ecological network to secondary extinctions, given a predefined species loss. Robustness is typically quantified as *R*_50_, which is the proportion of species one needs to remove such that 50% of all species goes extinct, after having accounted for secondary extinctions [8, 15]. Robustness is linked to the slope of the link-species distribution [4, 8, 13, 15], and so our results imply that environmental condition changes do not affect robustness. However, this does not imply that the absolute number of extinctions required to drive 50% of the species to extinction remains equal. For example, if a network has *R*_50_ = 0.4 but a million species, then 400, 000 extinctions are required to remove half of all species. As this network loses species and therefore gets smaller, however, fewer and fewer extinctions are needed to remove half of the species that are left. Thus, even if *R*_50_ remains constants along extinction trajectories, the impact of a single extinction on species loss gets larger.

We cannot be sure that modifications of the intercept of the log(*L*) ∼ log(*S*) relationship are biological phenomena or due to sampling biases. Indeed, detecting all the interactions within an ecosystem is challenging, and ecological network data can suffer from incomplete samplings [11, 13, 57–60]. As some environmental conditions (such as tree density) can affect sampling efficiency, it is thus possible that environmental condition changes have modified the intercept of the log(*L*) ∼ log(*S*) relationship [56,61] because links were harder to detect in general. For example, this can happen when environmental change makes populations smaller, and therefore more difficult to sample interactions between them [56, 59, 60, 62].

In conclusion, we studied ecological networks in ecosystems under various environmental conditions. We found link-species relationships that were ecosystem-specific but constant across most environmental conditions, despite profound changes of network size and structure. This result confirms the hypothesis proposed by Warren in 1990 [29], and is encouraging for conservation efforts. Indeed, it suggests that the impacts of species gain or loss transcend the underlying environmental drivers. The explanation of why some changes in environmental conditions did impact the link-species relationship is still poorly understood and subject of future research [10]. The parameters of this relationship have proposed to be linked to environmental conditions [18, 63], but methodological challenges remain [44]. Addressing these challenges is a next step to understanding how changes in environmental conditions affect ecosystems.

## Methods

Ecosystem interactions can be represented as ecological networks, where species are nodes and interactions (or links) are edges [4, 11–13, 64]. Ecological networks can be ob-served multiple times in the field, with each observation representing a part of the network. We searched the literature to collect ecological network datasets with at least eight observations per ecosystem (to allow for meaningful linear regressions), ensuring that these datasets were based on field observations rather than simulations. When datasets included multiple ecosystems, as indicated by their original authors, we treated each ecosystem as a separate entity. Network observations were coded as undirected adjacency matrices, where 0 and 1 indicated the absence or presence of an interaction, respectively. For each network observation, we recorded the number of species (*S*), the number of interactions (or links, *L*), and the composition of observed species and links. In cases where organisms were not identified at the species level, the term “species” referred to taxa.

Additionally, ecological network observations were derived from the same network sampled under varying environmental conditions. These conditions included environmental noise, which represents random variability between observations (e.g., spatial or temporal variability), and environmental gradients, which result from ecosystem manipulations such as different environmental contexts or human impacts (see supplementary material A).

To investigate the effects of changes in these environmental conditions on ecological network size, we performed mixed and non-mixed linear models for each ecosystem. The response variables were the number of species and the number of links (*L*) for each network observation. Environmental noise was treated as a random explanatory variable, while environmental gradients were treated as fixed explanatory variables. If the number of observations was fewer than six per environmental gradient level, all environmental conditions were treated as random explanatory variables. We selected the best models based on AIC scores and identified significant effects of explanatory variables (p-value *<* 0.05).

Furthermore, we assessed *β* diversity in species and links composition. For species *β* diversity, we calculated the Jaccard dissimilarity index, which can quantify the proportion of non-shared species between pairs of observations within the same ecosystem [42]. To assess links *β* diversity, we estimated rewiring rates, defined as changes in interaction composition among shared species between two observations. [5, 27, 28]. We used Novotny’s rewiring index to measure the ratio of unique interactions among shared species. However, since this ratio is influenced by the proportion of shared species (and thus the Jaccard index), we accounted for this dependency in our analysis [27, 43] (see supplementary material C).

Nevertheless, we examined the effects of ecological condition changes on the link-species relationship. To do this, we performed linear models with log(*L*) as the response variable and log(*S*) as a fixed explanatory variable. As in the network size models, environmental gradients were included as fixed variables (except when there were fewer than six observations per level), and environmental noise were included as a random variable. For each ecosystem, we selected the best model using AIC scores. We interpreted the effects of environmental conditions as changes in the intercept of the link-species relationship (Fig. 1 A) and effects of their interactions with log(*S*) as changes in the slope (Fig. 1 B). We also identified misaligned observations by locating points outside the 95% prediction interval. Finally, we estimated the *R*^2^ for each log(*L*) ∼ log(*S*) regression line.

## Supporting information

All supplementary material

## References

[1] Jacques Gignoux, Ian D. Davies, Shayne R. Flint, and Jean-Daniel Zucker. The Ecosystem in Practice: Interest and Problems of an Old Definition for Constructing Ecological Models. Ecosystems, 14(7):1039–1054, November 2011.

[2] William L. Geary, Michael Bode, Tim S. Doherty, Elizabeth A. Fulton, Dale G. Nimmo, Ayesha I. T. Tulloch, Vivitskaia J. D. Tulloch, and Euan G. Ritchie. A guide to ecosystem models and their environmental applications. Nature Ecology & Evolution, 4(11):1459–1471, November 2020.

[3] Geoffrey Caron-Lormier, David A. Bohan, Cathy Hawes, Alan Raybould, Alison J. Haughton, and Roger W. Humphry. How might we model an ecosystem? Ecological Modelling, 220(17):1935– 1949, September 2009.

[4] Pietro Landi, Henintsoa O. Minoarivelo, Åke Brännström, Cang Hui, and Ulf Dieckmann. Complexity and stability of ecological networks: A review of the theory. Population Ecology, 60(4):319–345, October 2018.

[5] Camila da Silva Goldas, Luciana Regina Podgaiski, Carolina Veronese Corrêa da Silva, Pedro Maria Abreu Ferreira, Jeferson Vizentin-Bugoni, and Milton de Souza Mendonça. Structural resilience and high interaction dissimilarity of plant–pollinator interaction networks in fire-prone grasslands. Oecologia, November 2021.

[6] Enrique Burgos, Horacio Ceva, Roberto P. J. Perazzo, Mariano Devoto, Diego Medan, Martín Zimmermann, and Ana María Delbue. Why nestedness in mutualistic networks? Journal of Theoretical Biology, 249(2):307–313, November 2007.

[7] Jordi Bascompte. Mutualistic networks. Frontiers in Ecology and the Environment, 7(8):429–436, 2009.

[8] Jennifer A. Dunne, Richard J. Williams, and Neo D. Martinez. Network structure and biodiversity loss in food webs: Robustness increases with connectance. Ecology Letters, 5(4):558–567, 2002.

[9] Jordi Bascompte and Daniel B. Stouffer. The assembly and disassembly of ecological networks. Philosophical Transactions of the Royal Society B: Biological Sciences, 364(1524):1781–1787, June 2009.

[10] David Moreno-Mateos, Antton Alberdi, Elly Morriën, Wim H. van der Putten, Asun Rodríguez-Uña, and Daniel Montoya. The long-term restoration of ecosystem complexity. Nature Ecology & Evolution, 4(5):676–685, May 2020.

[11] Tanya Strydom, Michael D. Catchen, Francis Banville, Dominique Caron, Gabriel Dansereau, Philippe Desjardins-Proulx, Norma R. Forero-Muñoz, Gracielle Higino, Benjamin Mercier, Andrew Gonzalez, Dominique Gravel, Laura Pollock, and Timothée Poisot. A roadmap towards predicting species interaction networks (across space and time). Philosophical Transactions of the Royal Society of London. Series B, Biological Sciences, 376(1837):20210063, November 2021.

[12] Thomas C. Ings, José M. Montoya, Jordi Bascompte, Nico Blüthgen, Lee Brown, Carsten F. Dormann, François Edwards, David Figueroa, Ute Jacob, J. Iwan Jones, Rasmus B. Lauridsen, Mark E. Ledger, Hannah M. Lewis, Jens M. Olesen, F.J. Frank Van Veen, Phil H. Warren, and Guy Woodward. Review: Ecological networks – beyond food webs. Journal of Animal Ecology, 78(1):253–269, 2009.

[13] Stephen R Proulx, Daniel E L Promislow, and Patrick C Phillips. Network think-ing in ecology and evolution. CellPress, page 9, 2005.

[14] Jian D. L. Yen, Reniel B. Cabral, Mauricio Cantor, Ian Hatton, Susanne Kortsch, Joana Patrício, and Masato Yamamichi. Linking structure and function in food webs: Maximization of different ecological functions generates distinct food web structures. Journal of Animal Ecology, 85(2):537–547, 2016.

[15] Camille Carpentier, György Barabás, Jürg Werner Spaak, and Frederik De Laender. Reinterpreting the relationship between number of species and number of links connects community structure and stability. Nature Ecology & Evolution, pages 1–8, May 2021.

[16] Ulrich Brose, Annette Ostling, Kateri Harrison, and Neo D. Martinez. Unified spatial scaling of species and their trophic interactions. Nature, 428(6979):167–171, March 2004.

[17] Anders Nielsen and Jordi Bascompte. Ecological Networks, Nestedness and Sampling Effort. Journal of Ecology, 95(5):1134–1141, 2007.

[18] Jean P. Gibert and Daniel J. Wieczynski. Constraints and variation in food web link-species space. Biology Letters, 17(4):20210109, April 2021.

[19] Neo D. Martinez. Constant Connectance in Community Food Webs. The American Naturalist, 139(6):1208–1218, June 1992.

[20] J. E. Cohen and F. Briand. Trophic links of community food webs. Proceedings of the National Academy of Sciences, 81(13):4105–4109, July 1984.

[21] R. Milo, S. Shen-Orr, S. Itzkovitz, N. Kashtan, D. Chklovskii, and U. Alon. Network Motifs: Simple Building Blocks of Complex Networks. Science, 298(5594):824–827, October 2002.

[22] Oscar Godoy, Fernando Soler-Toscano, José R. Portillo, and José A. Langa. The assembly and dynamics of ecological communities in an ever-changing world. Ecological Monographs, n/a(n/a):e1633, September 2024.

[23] Jason M. Tylianakis, Laura B. Martínez-García, Sarah J. Richardson, Duane A. Peltzer, and Ian A. Dickie. Symmetric assembly and disassembly processes in an ecological network. Ecology Letters, 21(6):896–904, 2018.

[24] Simon T. Segar, Tom M. Fayle, Diane S. Srivastava, Thomas M. Lewinsohn, Owen T. Lewis, Vojtech Novotny, Roger L. Kitching, and Sarah C. Maunsell. The Role of Evolution in Shaping Ecological Networks. Trends in Ecology & Evolution, 35(5):454–466, May 2020.

[25] Ulrich Brose, Julia L. Blanchard, Anna Eklöf, Nuria Galiana, Martin Hartvig, Myriam R. Hirt, Gregor Kalinkat, Marie C. Nordström, Eoin J. O’Gorman, Björn C. Rall, Florian D. Schneider, Elisa Thébault, and Ute Jacob. Predicting the consequences of species loss using size-structured biodiversity approaches. Biological Reviews, 92(2):684–697, 2017.

[26] Emma-Liina Marjakangas, Gabriel Muñoz, Shaun Turney, Jörg Albrecht, Eike Lena Neuschulz, Matthias Schleuning, and Jean-Philippe Lessard. Trait-based inference of ecological network assembly: A conceptual framework and methodological toolbox. Ecological Monographs, 92(2):e1502, 2022.

[27] Carmelo Gómez-Martínez and Amparo Lázaro. A new tool to improve the estimates of interaction rewiring considering the whole community composition. Methods in Ecology and Evolution, 15(8):1438–1449, 2024.

[28] Paolo Biella, Asma Akter, Jeff Ollerton, Anders Nielsen, and Jan Klecka. Anempirical attack tolerance test alters the structure and species richness of plant– pollinator networks. Functional Ecology, 34(11):2246–2258, 2020.

[29] Philip H. Warren. Variation in Food-Web Structure: The Determinants of Connectance. The American Naturalist, 136(5):689–700, 1990.

[30] Spencer A. Wood, Roly Russell, Dieta Hanson, Richard J. Williams, and Jennifer A. Dunne. Effects of spatial scale of sampling on food web structure. Ecology and Evolution, 5(17):3769–3782, September 2015.

[31] Leonardo A. Saravia, Tomás I. Marina, Nadiah P. Kristensen, Marleen De Troch, and Fernando R. Momo. Ecological network assembly: How the regional metaweb influences local food webs. Journal of Animal Ecology, 91(3):630– 642, 2022.

[32] Anders Nielsen and Ørjan Totland. Structural properties of mutualistic networks withstand habitat degradation while species functional roles might change. Oikos, 123(3):323–333, 2014.

[33] Jeanne David and Renate A. Wesselingh. Étude de la biodiversité des espaces verts gérés par l’UCLouvain sur le campus de Louvain-la-Neuve: type de gestion et réseaux plantes-pollinisateurs. PhD thesis, Catholic university of Louvain La Neuve, 2023.

[34] Moisès Guardiola, Constanti Stefanescu, Ferran Roda, and Joan Pino. Do asynchronies in extinction debt affect the structure of trophic networks? A case study of antagonistic butterfly larvae– plant networks - Guardiola - 2018 - Oikos - Wiley Online Library. Oikos, November 2017.

[35] Colin Olito and Jeremy W. Fox. Species traits and abundances predict metrics of plant–pollinator network structure, but not pairwise interactions. Oikos, 124(4):428–436, 2015.

[36] Julio M. Alcántara and Pedro J. Rey. Linking Topological Structure and Dynamics in Ecological Networks. The American Naturalist, August 2012.

[37] Chun-Huo Chiu, Anne Chao, Sebastian Vogel, Peter Kriegel, and Simon Thorn. Quantifying and estimating ecological network diversity based on incomplete sampling data. Philosophical Transactions of the Royal Society B: Biological Sciences, 378(1881):20220183, May 2023.

[38] Ellen A.R. Welti, Fan Qiu, Hannah M. Tetreault, Mark Ungener, John Blair, and Anthony Joern. Fire, grazing and climate shape plant–grasshopper interactions in a tallgrass prairie. Functional Ecology, December 2018.

[39] Els M. van der Zee, Christine Angelini, Laura L. Govers, Marjolijn J. A. Christianen, Andrew H. Altieri, Karin J. van der Reijden, Brian R. Silliman, Johan van de Koppel, Matthijs van der Geest, Jan A. van Gils, Henk W. van der Veer, Theunis Piersma, Peter C. de Ruiter, Han Olff, and Tjisse van der Heide. How habitat-modifying organisms structure the food web of two coastal ecosystems. Proceedings of the Royal Society B: Biological Sciences, 283(1826):20152326, March 2016.

[40] Jurene E. Kemp, Darren M. Evans, Willem J. Augustyn, and Allan G. Ellis. Invariant antagonistic network structure despite high spatial and temporal turnover of interactions. Ecography, 40(11):1315–1324, 2017.

[41] Serguei Saavedra, Simone Cenci, Ek Del-Kal, Karina Boege, and Rudolf P. Rohr. Reorganization of interaction networks modulates the persistence of species in late successional stages. Journal of Animal Ecology, May 2017.

[42] Paul Jaccard. Distribution comparée de la flore alpine dans quelques régions des Alpes orientales. Bulletin de la Murithienne, 1902.

[43] Vojtech Novotny. Beta-diversity of plant-insect food webs in tropical forests: A conceptual framework. Insect Conservation and Diversity, 2009.

[44] Chris Brimacombe, Korryn Bodner, Dominique Gravel, Shawn J. Leroux, Timothée Poisot, and Marie-Josée Fortin. Publication-driven consistency in food web structures: Implications for comparative ecology. Ecology, n/a(n/a):e4467, November 2024.

[45] Jose M. Montoya and Ricard V. Solé. Topological properties of food webs: From real data to community assembly models. Oikos, 102(3):614–622, 2003.

[46] Jens Olesen, Jordi Bascompte, Heidi Elberling, and Pedro Jordano. Temporal dynamics in a pollination network. Ecology, 89:1573–82, July 2008.

[47] X. Chen and J. E. Cohen. Support of the hyperbolic connectance hypothesis by qualitative stability of model food webs. Community Ecology, 1(2):215– 225, January 2006.

[48] Núria Galiana, Miguel Lurgi, Vinicius A. G. Bastazini, Jordi Bosch, Luciano Cagnolo, Kevin Cazelles, Bernat Claramunt-López, Carine Emer, Marie-Josée Fortin, Ingo Grass, Carlos Hernández-Castellano, Frank Jauker, Shawn J. Leroux, Kevin McCann, Anne M. McLeod, Daniel Montoya, Christian Mulder, Sergio Osorio-Canadas, Sara Reverté, Anselm Rodrigo, Ingolf Steffan-Dewenter, Anna Traveset, Sergi Valverde, Diego P. Vázquez, Spencer A. Wood, Dominique Gravel, Tomas Roslin, Wilfried Thuiller, and José M. Montoya. Ecological network complexity scales with area. Nature ecology & evolution, 6(3):307, January 2022.

[49] Salvador Sánchez-Carrillo, David G. Angeler, Miguel Álvarez-Cobelas, and Carmen Rojo. Abiotic drivers of consumer foodweb structure in lakes. Freshwater Science, 37(2):404–416, June 2018.

[50] Elisa Thébault and Colin Fontaine. Stability of ecological communities and the architecture of mutualistic and trophic networks. Science (New York, N.Y.), 329(5993):853–856, August 2010.

[51] Philip H. Warren. Estimating morphologically determined connectance and structure for food webs of freshwater invertebrates. Freshwater Biology, 33(2):213– 221, 1995.

[52] Jofre Carnicer, Pedro Jordano, and Carlos J. Melián. The temporal dynamics of resource use by frugivorous birds: A network approach. Ecological society of America, July 2009.

[53] Kirk O. Winemiller. Spatial and temporal variation in tropical fish trophic networks. Ecological Monographs, September 1990.

[54] Loïc Pellissier, Camille Albouy, Jordi Bascompte, Nina Farwig, Catherine Graham, Michel Loreau, Maria Alejandra Maglianesi, Carlos J. Melián, Camille Pitteloud, Tomas Roslin, Rudolf Rohr, Serguei Saavedra, Wilfried Thuiller, Guy Woodward, Niklaus E. Zimmermann, and Dominique Gravel. Comparing species interaction networks along environmental gradients. Biological Reviews, 93(2):785–800, 2018.

[55] Darren M. Evans, Michael J. O. Pocock, and Jane Memmott. The robustness of a network of ecological networks to habitat loss. Ecology Letters, 16(7):844–852, 2013.

[56] Jason M. Tylianakis, Etienne Laliberté, Anders Nielsen, and Jordi Bascompte. Conservation of species interaction networks. Biological Conservation, 143(10):2270–2279, October 2010.

[57] William Godsoe, Nathaniel J. Holland, Chris Cosner, Bruce E. Kendall, Angela Brett, Jill Jankowski, and Robert D. Holt. Interspecific interactions and range limits: Contrasts among interaction types. Theoretical Ecology, 10(2):167–179, June 2017.

[58] Pedro Jordano. Sampling networks of ecological interactions. Functional Ecology, 30(12):1883–1893, 2016.

[59] J. Christopher D. Terry and Owen T. Lewis. Finding missing links in interaction networks. Ecology, 101(7):e03047, 2020.

[60] Jens M. Olesen, Jordi Bascompte, Yoko L. Dupont, Heidi Elberling, Claus Rasmussen, and Pedro Jordano. Missing and forbidden links in mutualistic networks. Proceedings of the Royal Society B: Biological Sciences, 278(1706):725–732, September 2010.

[61] Etienne Laliberté and Jason M. Tylianakis. Deforestation homogenizes tropical parasitoid–host networks - Laliberté - 2010 - Ecology - Wiley Online Library. Ecological society of America, June 2010.

[62] Abhay Krishna, Paulo R. Guimarães Jr, Pedro Jordano, and Jordi Bascompte. A neutral-niche theory of nestedness in mutualistic networks. Oikos, 117(11):1609– 1618, 2008.

[63] Lydia Beaudrot, Miguel A. Acevedo, Daniel Gorczynski, and Nyeema C. Harris. Geographic differences in body size distributions underlie food web connectance of tropical forest mammals. Scientific Reports, 14(1):6965, March 2024.

[64] Ellie Nagaishi and Kazuhiro Takemoto. Network resilience of mutualistic ecosystems and environmental changes: An empirical study. Royal Society Open Science, 5(9):180706, September 2018.

