## Supplementary material for "Network theory predicts ecosystem robustness across environmental conditions": All supplementary material

#### A Data summary

| Author | Name | N ecosystem | Environmental gradient | Symbol | N observations |
|---|---|---|---|---|---|
| Alcantara *et al.* | Plant post fire | 1 | Recent fire<br>Ancient fire | ●<br>● | 5<br>5 |
| Chiu *et al.* | Tree beetles | 1 | Open environment<br>Naturally shaded<br>Artificially shaded | ■<br>▲<br>● | 24<br>24<br>24 |
| David *et al.* | City pollination | 1 | Grass reaping<br>Grass cutting | ▲<br>● | 54<br>66 |
| Guardiola *et al.* | Butterfly predation | 1 | | ● | 26 |
| Kemp *et al.* | Insects predation | 1 | Spring<br>Autumn | ▲<br>● | 6<br>6 |
| Nielsen *et al.* | Forest gestion | 1 | Clear cut<br>Young forest<br>Old growth forest | ●<br>●<br>● | 4<br>4<br>4 |
| Olito *et al.* | Mountain pollination | 1 | Altitude | ●▲■ | 80 |
| Saavedra *et al.* | Butterfly succession | 1 | Middle succession state<br>Late succession state<br>Initial succession state | ■<br>▲<br>● | 12<br>12<br>12 |
| Saravia *et al.* | Several marine ecosystems | 6 | | ● | 254 |
| Van Der Zee *et al.* | Several coastal ecosystems | 2 | Colonized soil<br>Cobblestone soil<br>Bare soil | ■<br>▲<br>● | 8<br>8<br>8 |
| Welti *et al.* | Grasshopper predation | 1 | 4 fire/year & no buffalos<br>4 fire/year & buffalos<br>20 fire/year & no buffalos<br>20 fire/year & buffalos<br>1 fire/years & no buffalos<br>1 fire/years & buffalos | $*$<br>⊠<br>+<br>■<br>▲<br>● | 19<br>13<br>19<br>13<br>19<br>13 |
| Wood *et al.* | Alaska islands | 1 | | ● | 348 |

Summary of each dataset with original author (**Author**), ecosystem(s) name (**Name**), number of ecosystems for an author (**N ecosystem**), environmental gradient levels (**Environmental gradient**), its symbol (**Symbol**) for figures 2, 4 and 5 and number of observations in each gradient (**N observations**).

#### B Detailled results

| Ecosystem | Condition | Gradient/noise | Species | Links | Intercept | Slope |
|---|---|---|---|---|---|---|
| Plant post fire | Last fire | Gradient | NS | NS | NS | NS |
| Tree beetles | Sun exposure | Gradient | ** | *** | *** | NS |
| | Time | Noise | . | ** | NS | NS |
| | Plot | Noise | NS | NS | NS | NS |
| City pollination | Grass mowing | Gradient | *** | *** | . | NS |
| | Time | Noise | NS | NS | NS | NS |
| | Plot | Noise | NS | NS | NS | NS |
| Insects predation | Season | Gradient | NS | NS | NS | NS |
| Forest gestion | Gestion | Gradient | NS | NS | NS | NS |
| Mountain pollination | Altitude | Gradient | . | NS | . | NS |
| | Time | Noise | *** | *** | NS | NS |
| Butterfly succession | Succession state | Gradient | NS | * | NS | NS |
| | Time | Noise | ** | ** | NS | NS |
| | Plot | Noise | NS | NS | NS | NS |
| Several coastal ecosystems | Soil composition | Gradient | *** | *** | NS | NS |
| | Plot | Noise | NS | NS | NS | NS |
| Grasshopper predation | Buffalo | Gradient | *** | *** | * | NS |
| | Fire frequency | Gradient | ** | NS | * | NS |
| | Buffalo:fire | Gradient | ** | . | NS | NS |
| | Time | Noise | *** | *** | . | NS |

Summary of impacts from changes in environmental conditions (**Condition** that are gradient or noise according to **Gradient/noise**) on number of species (**Species**), links (**Links**), $\log(L) \sim \log(S)$ intercept (**Intercept**) and slope (**Slope**) in each ecosystem (**Ecosystem**). Symbols are relative to p-values, *** is for p-values inferior to $0.001$, ** for p-values between $0.001$ and $0.01$, * for p-values between $0.01$ and $0.05$, . for p-values between $0.05$ and $0.1$, and NS for p-values superior to $0.1$. Only *, **, and *** were interpreted as significant.

#### C Index equations

The Jaccard dissimilarity index quantifies the dissimilarity between two groups by measuring the ratio between shared individuals and all individuals in the following equation:

$$J = 1 - \frac{A \cap B}{A \cup B}$$

With $A$ as the individuals of the first group and $B$ as the individuals of the second group. The final result is between $0$ (all individuals are shared) and $1$ (no individuals are shared). This index only considers the presence or absence of species [1]. Here we used it with ecological network observation as groups and species as individuals.

Between two ecological network observations, interactions can differ due to a modification of the species composition (a new species brings new interactions), or by rewiring (modification interactions between shared species) [2]. To estimate rewiring, we used Novotny's method which consists of making sub-networks of shared species and counting the number of unique interactions in them with the following equation:

$$\beta_{OS} = \frac{b_{OS} + c_{OS}}{2 \cdot a + b + c}$$

With $b_{OS}$ as the number of unique interactions of the first sub-network of shared species, $c_{OS}$ as the number of unique interactions of the second sub-network, $a$ as the number of shared interactions, $b$ as the number of unique interactions of the first network, and $c$ as the number of unique interactions of the second network. This index gives a result between $0$ (no rewiring or no common species) and $1$ (all species are shared but no interactions are shared) [3]. However, this index also depend of the proportion of shared species and so, have to be considered with species $\beta$ diversity index, such as the Jaccard dissimilarity index [2].

### D Jaccard plot

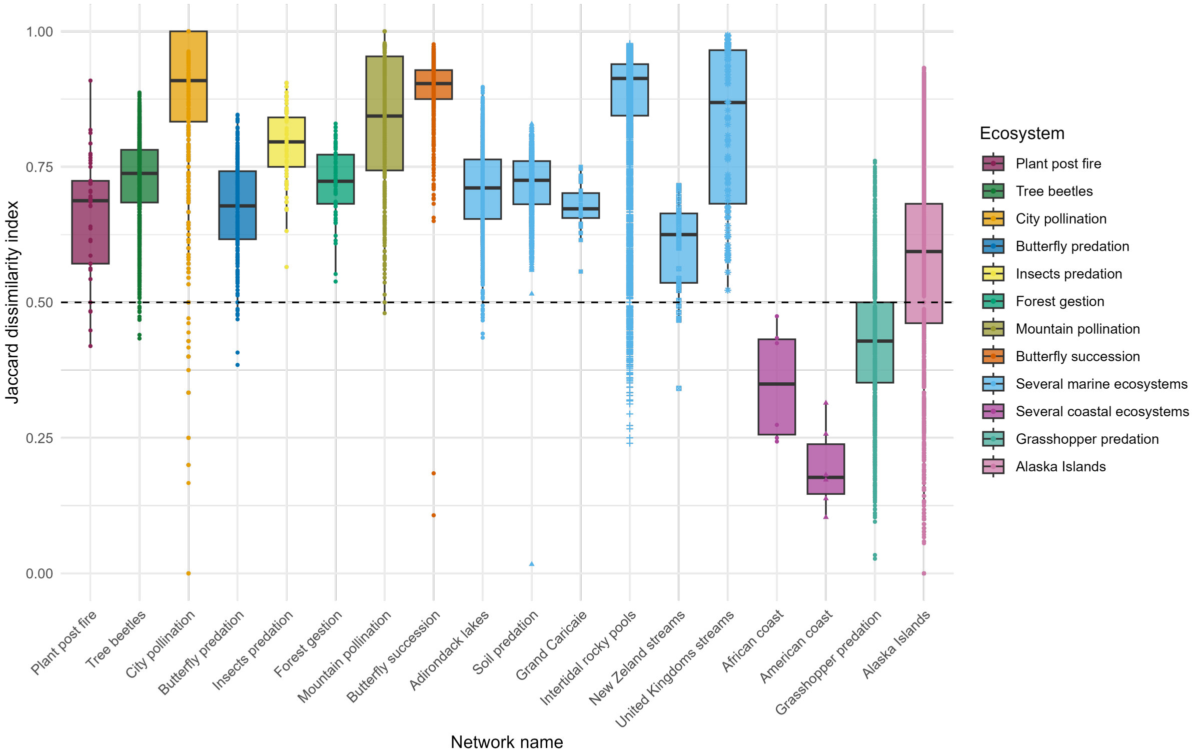


Jaccard dissimilarity index of species between every pair of observations of the same network. The dashed line refers to $J = 0.5$, so $50\%$ of shared species between the two observations.

### References


[1] Paul Jaccard. Distribution comparée de la flore alpine dans quelques régions des Alpes orientales. *Bulletin de la Murithienne*, 1902.

[2] Carmelo Gómez-Martínez and Amparo Lázaro. A new tool to improve the estimates of interaction rewiring considering the whole community composition. *Methods in Ecology and Evolution*, 15(8):1438–1449, 2024.

[3] Vojtech Novotny. Beta-diversity of plant-insect food webs in tropical forests: A conceptual framework. *Insect Conservation and Diversity*, 2009.
